# The Emergence of Circadian Timekeeping in the Intestine

**DOI:** 10.1101/2023.06.06.543952

**Authors:** Kathyani Parasram, Amy Zuccato, Minjeong Shin, Reegan Willms, Brian Deveale, Edan Foley, Phillip Karpowicz

## Abstract

The circadian clock is a molecular timekeeper, present from cyanobacteria to mammals, that coordinates internal physiology with the external environment. The clock has a 24-hour period however development proceeds with its own timing, raising the question of how these interact. Using the intestine of *Drosophila melanogaster* as a model for organ development, we track how and when the circadian clock emerges in specific cell types. We find that the circadian clock begins abruptly in the adult intestine and gradually synchronizes to the environment after intestinal development is complete. This delayed start occurs because individual cells at earlier stages lack the complete circadian clock gene network. As the intestine develops, the circadian clock is first consolidated in intestinal stem cells with changes in ecdysone and Bursicon hormone signalling influencing the transcriptional activity of Clk/cyc to drive the expression of *tim*, *Pdp1,* and *vri*. In the mature intestine, stem cell lineage commitment transiently disrupts clock activity in differentiating progeny, mirroring early developmental clock-less transitions. Our data show that clock function and differentiation are incompatible and provide a paradigm for studying circadian clocks in development and stem cell lineages.

## Introduction

Circadian rhythms are 24-hour cycles of physiological activity driven by the circadian clock, a molecular pacemaker found in nearly all cells of the body (Hardin 2011; Cox and Takahashi 2019). The circadian clock promotes health by coordinating tissues in the body to anticipate daily environmental changes (Panda 2016). In many animals, the circadian system is hierarchical, consisting of a central pacemaker in the brain and peripheral pacemakers located in other organs (Reppert and Weaver 2002). How the circadian clock develops is not well understood, but it arises during early embryogenesis in fish (Dekens and Whitmore 2008) and late fetal and post-natal development in humans (Rivkees 2003). The central pacemaker in mice arises earlier than clocks in the rest of the body (Inada et al. 2014; Polidarová et al. 2014; Carmona-Alcocer et al. 2018) which suggests that certain tissue clocks are suppressed during development (Umemura and Yagita 2020). In *Drosophila melanogaster*, larval behaviours such as photo-avoidance show circadian rhythms (Mazzoni et al. 2005; Keene et al. 2011; Baik et al. 2017; Baik et al. 2018; Asirim et al. 2020) and the central pacemaker can be synchronized to the environment as early as first instar larva (Sehgal et al. 1992). These observations raise questions about when and how the circadian clock emerges in specific tissues, and the genetic and/or cellular mechanisms that underlie the birth of the timing mechanism.

The circadian clock of *Drosophila* is a transcription-translation cycle in which the Clock/cycle (Clk/cyc) heterodimer activates gene expression while two of its targets, *period* (*per*) (Hao et al. 1997) and *timeless* (*tim*) (McDonald et al. 2001), repress Clk/cyc activity to restart the cycle (Figure 1A) (Darlington et al. 1998). A secondary loop consisting of *Par-domain protein 1* (*Pdp1*) and *vrille* (*vri*) stabilizes these rhythms by regulating *Clk* (Cyran et al. 2003; Glossop et al. 2003; Hardin 2011). In *Drosophila* peripheral clocks can be synchronized to the environment indirectly by the central pacemaker in the brain (Hardin 2011) and directly through the blue-light photoreceptor *cryptochrome* (*cry*) (Emery et al. 2000).

**Figure 1:**
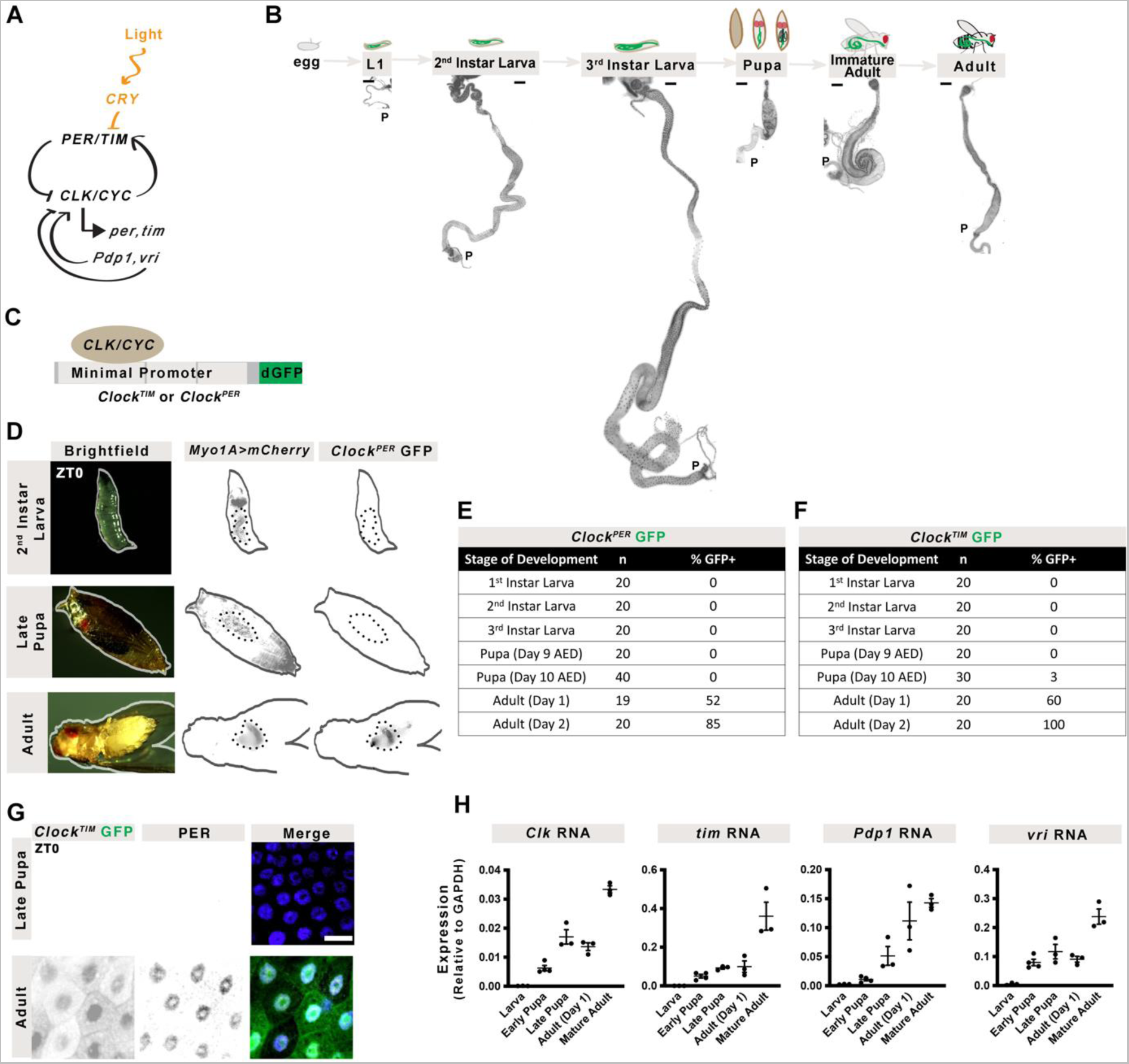
The Circadian Clock Emerges in the Adult Intestine. **(A)** The circadian clock comprises the transcriptional activators Clk/cyc and their repressors per/tim. The repressor vri and activator Pdp1 form a secondary feedback system and light is sensed through cry. **(B)** *Drosophila* development from egg through three larval instars, followed by pupation prior to eclosion as an adult. Images of the midgut from larva to adults shows dramatic growth and remodelling of the intestine, DAPI stains nuclei (black). All images shown at same scale (scale bar = 200µm). “P” indicates posterior. (**C**) Schematic of the Clock Reporter showing binding of Clk/cyc to the minimal promoter of either *tim* (for the *Clock^TIM^* reporter) or *per* (*Clock^PER^*) located upstream of destabilized GFP. **(D)** Images of larva, pupa, and adult show that *Clock^PER^* is expressed only in adult intestines (*Myo1A>mCherry*). Quantification of the number of flies expressing **(D)** *Clock^PER^* or **(E)** *Clock^TIM^* in the intestine throughout development. **(G)** Antibody staining showing that PER protein is only expressed in the adults and is localized to the nuclei in *Clock^TIM^* flies at ZT0 (Zeitgeber Time relative to photoperiod, ZT0=lights on, ZT12=lights off). Scale 10µm. Histone marks nuclei. **(H)** The mRNA expression of circadian clock genes at ZT0 from 3^rd^ instar larva to adults showing the circadian clock transcripts increase in adults. Error bars indicate ±SEM. One-Way ANOVA. Each point represents one replicate (20 intestines). Full statistics are shown in Supplemental Information. Related to S1-2.

To address the question of how circadian clock activity is established during development, we tested a tissue with a simple lineage (Micchelli and Perrimon 2006; Ohlstein and Spradling 2006), well-established developmental stages (Tepass and Hartenstein 1994; Takashima et al. 2011b), and adult circadian clock activity (Karpowicz et al. 2013; Parasram et al. 2018). The adult intestine is derived from a population of cells known as adult midgut precursors (AMPs) that proliferate throughout larval development (Hartenstein et al. 1992; Hartenstein and Jan 1992; Jiang and Edgar 2009; Mathur et al. 2010) and pupation (Mathur et al. 2010; Micchelli et al. 2011; Takashima et al. 2011a), until the intestine matures three days after emergence as an adult (eclosion) (Hartenstein and Jan 1992; Jiang and Edgar 2009; Buchon et al. 2013). The intestine of *Drosophila* consists of intestinal stem cells (ISCs) which divide into progenitor cells, known as enteroblasts (EBs) or enteroendocrine mother cells, that then differentiate into enterocytes (ECs) or enteroendocrine cells (EEs), respectively (Micchelli and Perrimon 2006; Ohlstein and Spradling 2006; He et al. 2018). In the adult, the circadian clock controls ISC mitosis during regeneration (Karpowicz et al. 2013), and we have previously shown that epithelial cells of the intestine have clock function (Parasram et al. 2018).

Our results demonstrate that clock activity arises only after eclosion where it is synchronized to the environment by photoperiod and feeding over the first three days of adulthood. We characterize the transcriptome of single cells in the *Drosophila* intestine during adult maturation and clock formation and identify key regulators of this developmental transition. We propose a model where the ISCs are the first cells to have robust clock gene expression, that is disrupted during cellular differentiation to be resumed in specialized tissue cells. We further suggest that complete clock gene networks are absent in EEs and EBs, which express only a subset of clock genes. Insects represent the most abundant and diverse species of animal on Earth, hence our observations are significant in understanding how a critical physiological process is established in these animals and functions in their adulthood. The maturation of the *Drosophila* intestinal clock is also reminiscent of mammalian / human development, where tissue clocks are synchronized to daily timing postnatally. Thus, our results establish a paradigm for the birth of circadian timekeeping in animal tissues by providing a transcriptomic framework of clock development at the single cell level.

## Results

### The Circadian Clock Emerges in the Adult Intestine

During post-embryonic development the intestine undergoes significant growth and remodelling (Figure 1B). We first determined if the circadian clock affects intestinal development by comparing isogenic wildtype and *per^01^* mutant flies that exhibit no clock rhythms. Intestinal size, cell density, and the number of cells in AMP / PC islets do not differ significantly in *per^01^* mutants (Figure S1A-B), indicating that intestinal development is not affected in the absence of circadian clock activity. To determine when Clk/cyc transcriptional activity begins, we used two different reporter constructs (*Clock^PER^*and *Clock^TIM^*) inserted on different chromosomes that use destabilized GFP (dGFP) to visualize Clk/cyc transcriptional activity on *per* and *tim* promoters with ~1h temporal resolution (Parasram et al. 2018) (Figure 1C). The intestine was visualized using the GAL4/UAS system (*Myo1A>mCherry*) through the marker *Myosin 31DF* (*Myo1A)* that is expressed in both larval and adult intestinal cells (Morgan et al. 1995). No Clk/cyc transcriptional activity is present in larval (L1, L2, L3) or pupal flies, however, immediately after eclosion (adult day 1), Clk/cyc become active and remain so in later adulthood (Figure 1D-F) irrespective of reproduction status (Figure S1C-G. To verify our reporter, we probed for *per* protein using both an antibody and a C-terminal GFP fusion reporter (*per-AID-eGFP)* (Chen et al. 2018), and *cry* protein using an N-terminal fusion protein for *cry* (*cry-GFP*) (Agrawal et al. 2017). In all cases, these clock proteins are absent in pupa but present in recently-eclosed adults (Figure 1G, S2A-C). These results indicate that circadian clock activity begins only after pupal metamorphosis in the *Drosophila* intestine.

We next used RT-qPCR to examine the transcriptional expression of the core clock genes (*Clk, cyc, per, tim*), the secondary stabilizing loop (*Pdp1, vri*), the blue-light photoreceptor (*cry*), and the phosphatase *dbt*, a kinase that regulates *per* to reset Clk/cyc driven transcription (Kloss et al. 1998; Price et al. 1998). The intestinal expression of *Clk*, *tim*, *Pdp1E*, and *vri* increase throughout development to peak in adulthood, whereas *dbt*, *cyc*, *per*, and *cry* expression increase earlier in development before the stage when Clk/cyc activity exist (Figure 1H, S2D). Therefore, we hypothesized that the changes in clock activity are likely due to changes in *Clk, tim, Pdp1* and/or *vri*.

When we looked more closely at Clk/cyc activity in the last stages of pupation we noticed few scattered GFP cells (Figure S2E), raising the question of which intestinal cell type expresses Clk/cyc activity first. We measured Clk/cyc activity in pupal and adult ISC/EBs (marked by *esg>mCherry*) and ECs (marked by *Myo1A>mCherry*) and found that these early ISC/EBs show similar Clk/cyc activity in the late pupa and adults, unlike ECs that have significantly higher Clk/cyc activity in adults, suggesting that the ISC/EBs establish their circadian clock first (Figure S2F,G). Moreover, when the circadian clock is absent in all cells except ISC/EBs (*cyc^0^*clock-dead mutant with *esg>cyc* rescue) clock activity persists in the ISC/EBs indicating that these cells can develop clocks independently of the other clocks in the body (Figure S2H).

### Daily Clock Rhythms Are Established After Three Days of Adulthood

Since the intestinal clock emerges after pupation, we asked whether it is rhythmic immediately, with daily maxima and minima that are characteristic of circadian rhythms, or whether it needs time to synchronize to the environment. We tested wildtype *CantonS* fly intestines by RT-qPCR to compare adult day 1 (immature) and day 4 (mature) gene expression patterns over a 24-hour period. On day 1, the transcripts of most clock components are not rhythmic suggesting the transcriptional cycle is not yet present. Genes such as *Pdp1* then increase in rhythmicity three days later, and *cry* completely changes the timing of its initial rhythms (Figure 2A). Of note, the expression of all clock genes, with the exception of *cry,* are lower on day 1 than day 4, suggesting that establishment of robust transcriptional cycles takes several days.

**Figure 2:**
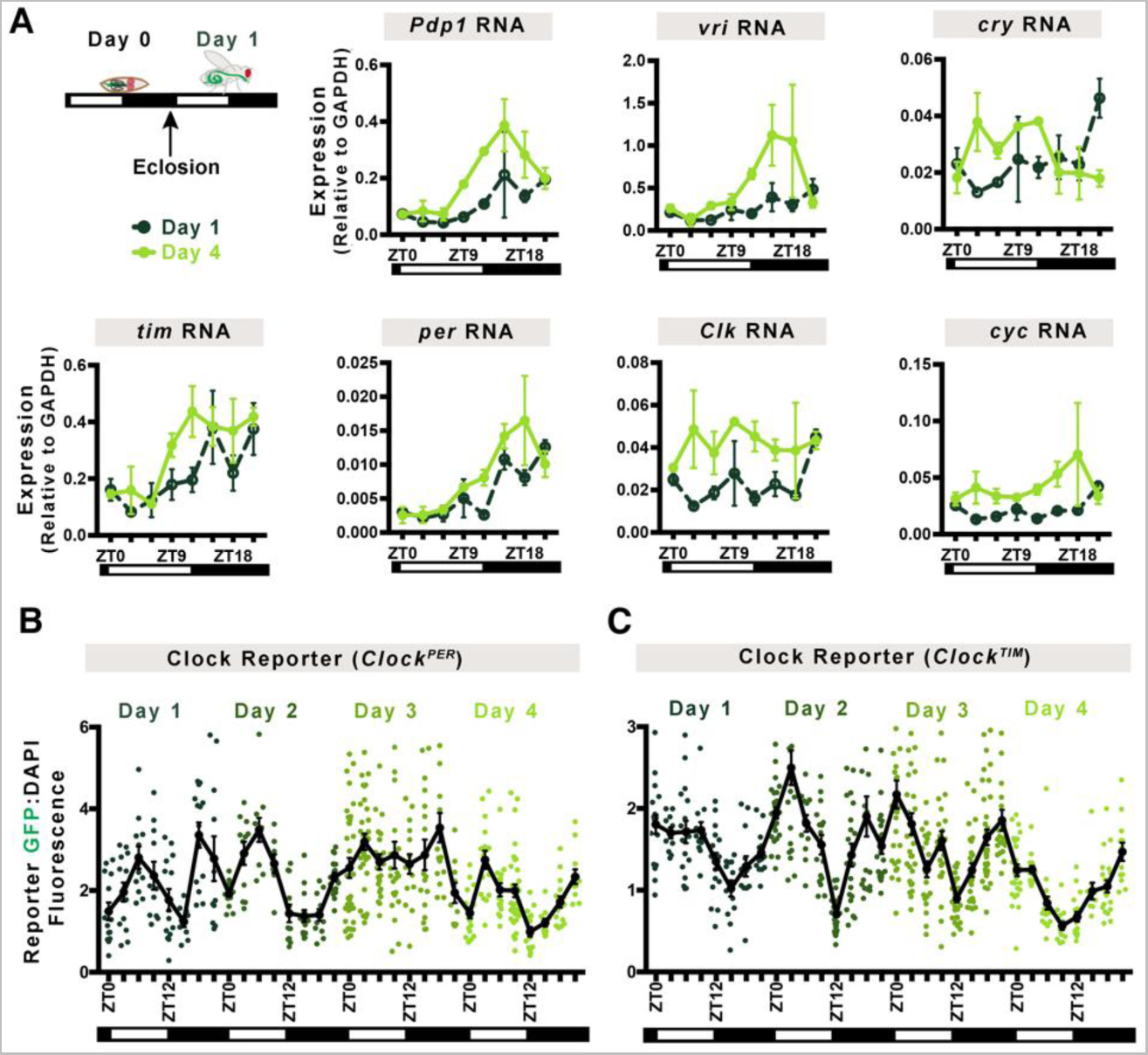
Daily Rhythms Are Established After Three Days of Adulthood. **(A)** Circadian clock genes are expressed in adults immediately after eclosion (day 1, shown in the schematic) but rhythms are stronger at day 4. For instance, amplitude of mRNA rhythm of *Pdp1* increases, and *cry* timing changes, other genes similarly show differences as intestine matures. Line shows mean. Error bars indicate ±SEM. Each point is the average of two replicates each consisting of 15-20 intestines. Two-Way ANOVA. **(B)** Clk/cyc transcriptional activity is initially stochastic and shows mature rhythms only by day 4 for *Clock^PER^*with a peak around ZT0 and trough around ZT12. **(C)** Similar results are observed for *Clock^TIM^*indicating that both per and tim transcription are similarly regulated over development. Lines show mean. Error bars indicate ±SEM. Each point represents one intestine. Day 1 corresponds to the first light and dark cue after eclosion. One-Way ANOVA. Full statistics are shown in Supplemental Information. Related to S2-3.

The gradual emergence of transcriptional rhythmicity suggests Clk/cyc transactivation is not synchronized until the intestine matures. To test this, we imaged the *Clock^PER^* and *Clock^TIM^* reporters in the intestine following pupation every 3-hours over the first four days of adulthood (Figure 2B-C, S3A). When the adult ecloses, Clk/cyc transcriptional activity is present, but its rhythms are not 24-hour circadian oscillations until day 4 when these consolidate to exhibit a maximum in the morning (ZT21-ZT0) and minimum in the evening (ZT9-ZT12) characteristic of the *Drosophila* intestine (Parasram et al. 2018). This suggests Clk/cyc activity at both *per* (Figure 2B) and *tim* (Figure 2C) promoters are fully developed in the mature adult intestine. A defining characteristic of circadian rhythms is their ability to continue in the absence of environmental cues such as photoperiod (Bruce and Pittendrigh 1957). Accordingly, *Clock^TIM^* intestines were examined when flies were shifted to constant darkness (DD) on day 4-5 (Figure S3B). The rhythms are similar to those under LD conditions, with a maximum in the morning (CT21-0) and minimum in the evening (CT12), demonstrating that Clk/cyc activity is free-running at day 4. Taken together, our results show that circadian clock development in the intestine has three distinct phases: (1) embryogenesis to late pupa when the circadian clock circuit is absent, (2) between adult day 1 to 3 when the Clk/cyc are active but not yet synchronized to the environment, and (3) day 4 when the circadian clock is active and rhythmic in the mature intestine.

### Clock Gene Expression Increases in ISCs and ECs During Intestinal Maturation

Previous studies have not been able to examine circadian clock development in individual cells of a tissue, hence it is not clear if clock components mature homogenously, with individual cells gradually expressing all clock genes, or heterogeneously, with individual cells expressing certain clock genes but not others. To further test the development of the circadian pacemaker, we profiled transcriptional changes in *CantonS* intestinal development over each of the three stages in clock development, from early pupa, immature adults (day 1), and mature adults (day 7-8) using scRNA-seq. Early pupal intestines do not have Clk/cyc activity, immature adult intestines have Clk/cyc activity but no rhythms, and mature adult intestines have rhythmic Clk/cyc activity; we hypothesized that clock development would increase homogenously in all intestinal cell types over these developmental stages. We recovered 5190 cells (2013 pupal, 1197 immature adult, and 1980 adult cells) that show 23 different cell states (Figure 3A, S4A-B), consistent with previous scRNA-seq studies of the *Drosophila* intestine (Buchon et al. 2013; Marianes and Spradling 2013; Dutta et al. 2015; Hung et al. 2020b; Li et al. 2022), capturing the changing intestinal transcriptome throughout development. Our data provides a new resource of cellular development in the insect intestine.

**Figure 3:**
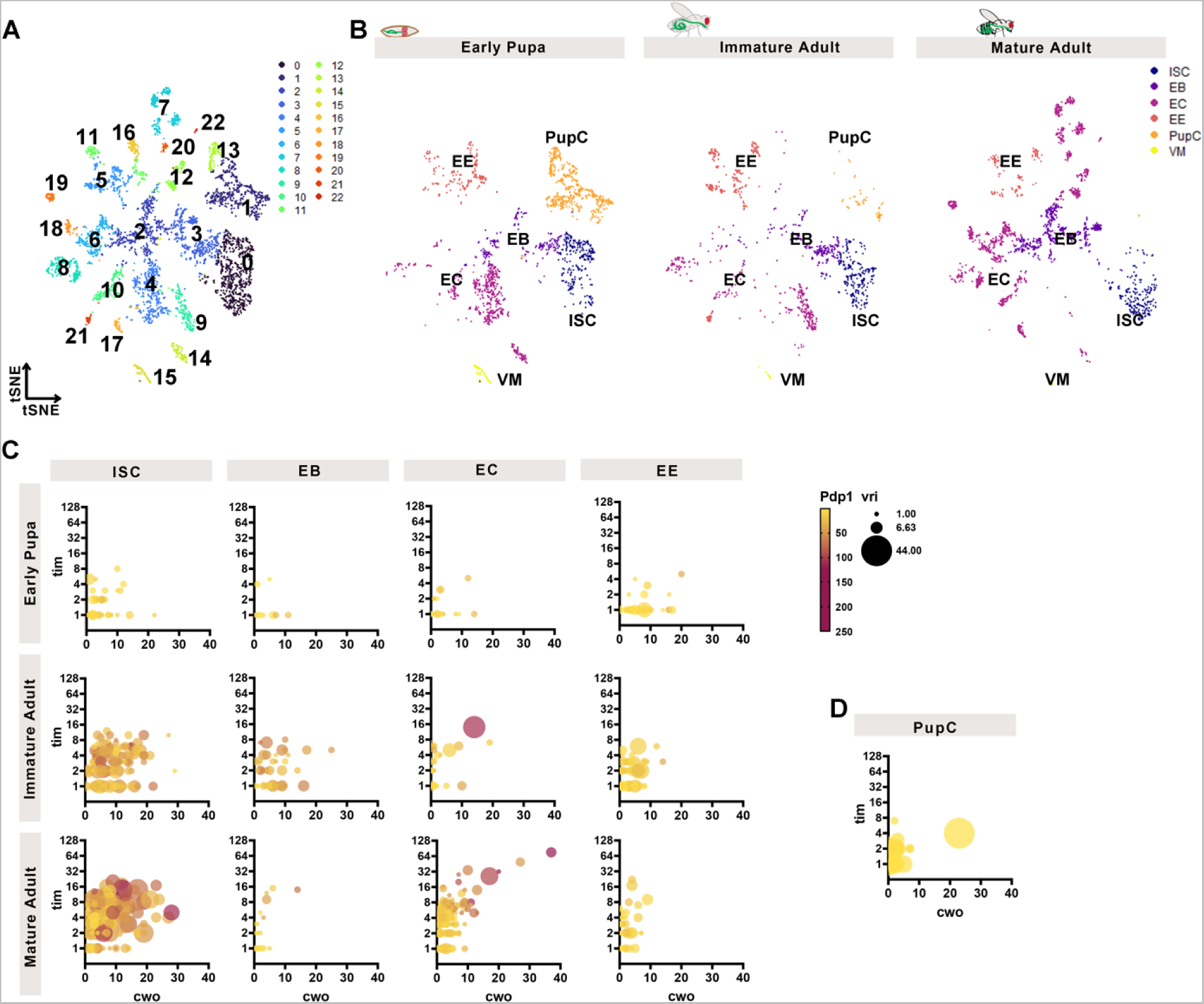
Clock Gene Expression Increases in ISCs and ECs During Intestinal Maturation. scRNA-seq analysis of clock gene expression of the intestine of early pupa, day 1 immature adults, and day 7-8 mature adults showing **(A)** tSNE plots with 23 different cell populations over the integrated dataset. **(B)** Population changes are evident during three developmental stages with PupCs (orange) disappearing from the pupal intestine, and ECs (purple) appearing in distinct clusters in the mature intestine. ISCs (blue) are present at all stages, similar to EEs (red). For cluster assignments and markers see Figure S4A, Table S10. **(C)** Multidimensional plots showing the changes in four highly expressed clock genes from early pupa to adult: *tim* (y-axis), *cwo* (x-axis), *Pdp1* (color), *vri* (size of datapoint). Initially expression of all four genes in ISCs (first column of graphs) is low but increases during maturation. This pattern is also seen in ECs (third column). EBs and EEs do not show these increases, cells remain clustered together at low levels of expression of all four genes. **(D)** PupCs also show low clock gene expression. See Supplemental Information for correlation matrices of multidimensional plots. Related to S4, Table S7-10.

During the transition from pupa to mature adult, the cell population of the *Drosophila* intestine undergoes cellular changes (Figure 3B). The ISCs and EEs are present at all stages but ECs are more restricted to later stages. A population of cells we denote as Pupal Cells (PupC) decrease in the immature adult and are completely absent in the mature adult. PupCs express some genes found in the EEs and ECs of the adult (Table S10, (Hung et al. 2020a)) and are likely derived from AMPs but represent a separate cell type only present in pupae (Figure S4C), consistent with previous reports that show transient pupal midgut cells disappear during metamorphosis (Takashima et al. 2011b). Several subtypes of differentiated ECs are not present in the pupae or immature adult and are only present in the mature adult, consistent with the maturation and growth of the *Drosophila* intestine (O’Brien et al. 2011). To assess circadian gene expression in these cells, we surveyed gene counts in individual cells (Figure S4D, Table S7-9). We were able to detect all core clock components (Figure S4D) whose expression at the population level increases as pupa transition to mature adults consistent with the RT-qPCR data (Figure 1H, S2D). We detected the genes *per, Clk*, and *cyc* at much lower levels than *tim, Pdp1, vri, dbt,* and *cwo*, a repressor of Clk/cyc. We therefore used the higher-expressed clock genes as a readout of clock development.

Previously, we reported that ISCs, EBs, and ECs, but not EEs have Clk/cyc transcriptional rhythms (Parasram et al. 2018). We therefore asked whether individual cells, including EEs, during these developmental stages express certain clock genes, but not the complete clock system. To test this, we focused on clock target genes that are more highly expressed and also feedback to regulate the core clock (*cwo, tim, Pdp1* and *vri*). We graphed all four genes in a multi-dimensional fashion to simultaneously determine changing expression of all four in individual cells (*tim* is y-axis, *cwo* is x-axis, *vri* is size, *Pdp1* is color). Individual ISCs show low expression of all four genes in the pupa, increasing co-expression in the immature adults, that further increases in the mature adults (Figure 3C). This suggests that the circadian clock program is not expressed in individual ISCs homogenously in the pupa, and that the arrhythmic Clk/cyc activity on day 1 is, at least in part, also due to low expression of the circadian clock program in individual cells. Individual ECs show very similar patterns of change, gradually single cells develop that express all four clock genes at higher levels. In contrast, clock genes are not expressed in individual EBs and EEs in the same manner. First, these do not show the robust expression of multiple clock genes at any developmental stage including the mature intestine. For example, only 10% of mature adult EBs coexpress 3-4 of clock genes compared to 98% of mature adult ISCs. Second, in many cells one or two clock genes are high (Figure S4C), but the remainder are low, suggesting clock components may have non-clock functions in specific subtypes of EEs or EBs but are not co-expressed as part of a circadian expression programme. The highest clock gene expression is present in ISCs, that increase expression in the immature and mature adult intestine, whereas the other intestinal cell types showed lower expression levels. Finally, individual PupCs do not express clock genes or express few at very low levels, which is consistent with the absence of Clk/cyc activity in the pupal intestine (Figure 3D). We conclude that ISCs and ECs increase clock gene expression after pupation, supporting a model where clock gene expression that is absent earlier in development increases in single cells homogenously to reach its maximum in the mature adult intestine.

### Hormonal Regulators of Clock Emergence

The sudden emergence of Clk/cyc activity in the immature adult intestine (Figure 1C-G) suggests that either a regulatory factor initiates the expression of circadian clock genes in the adult, or else suppresses them in the pupa, until the appropriate developmental stage is completed. Hormone nuclear receptors control many aspects of *Drosophila* physiology and metamorphosis (King-Jones and Thummel 2005; Tennessen and Thummel 2011) and are intimately connected with circadian clock function (Zhao et al. 2014). We therefore hypothesized that clock development is coordinated by hormone nuclear receptor signaling in the developing intestine. We used SCENIC (Aibar et al. 2017), a bioinformatic approach, that evaluates scRNA-seq transcriptomes to identify active gene networks and infer the underlying transcriptional regulators, to analyze pupal, immature, and mature adult cells. Identification of cell-specific transcription factors revealed enrichment of *klu* and *Sox21a* in ISCs, known to positively regulate ISC function (Meng and Biteau 2015; Hung et al. 2020a), the Notch target *E(spl)*) in ISC/EBs, known to drive ISC differentiation (Ohlstein and Spradling 2007; Guo and Ohlstein 2015), *nub*/*Pdm1* in ECs, a well-known marker of this cell type (Micchelli and Perrimon 2006; Ohlstein and Spradling 2006), *tap* in EEs (Guo et al. 2019), known to promote EE cell fate (Li et al. 2019) (Figure 4A, S4C). Hence SCENIC is able to identify cell specific transcriptional networks in the *Drosophila* intestine accurately.

**Figure 4:**
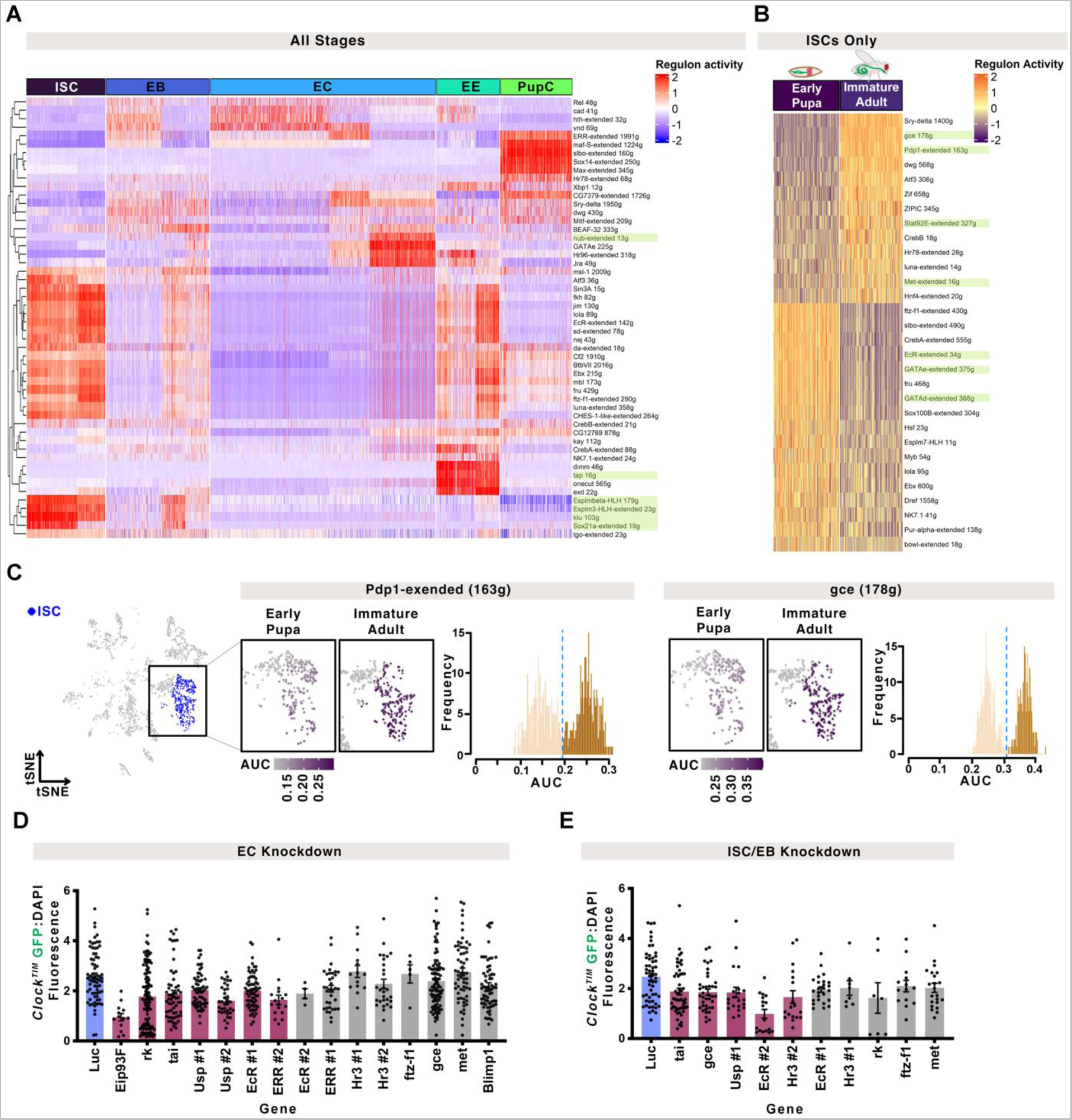
Hormonal Regulators of Clock Emergence. **(A)** SCENIC analysis of differentially expressed regulons that distinguish individual cell populations show clear changes in the transcriptional programs of different intestinal cell types (klu and Sox21a in ISCs, nub in ECs, tap in EEs). **(B)** Differential regulon expression focusing on the pupal-to-adult transition in ISCs shows that specific transcriptional pathways change during ISC development (*gce/met*, *Pdp1*, *Stat92E* are increased; *EcR*, *GATA* are decreased). Scaled AUC values are shown, ##g indicates the number of genes enriched in each regulon’s network. **(C)** The tSNE plot for early pupa and immature adult cells highlighting the ISC cluster used for SCENIC analysis, AUC values shown on a tSNE and in histograms showing population distributions based on regulon expression for the nuclear receptors *gce* and the clock gene *Pdp1*. **(D)** Knockdown of nuclear receptors in ECs or **(E)** ISC/EBs show that loss of hormone signalling transduction decreases Clk/cyc activity. One-way ANOVA., multiple comparison, p-value<0.05 (shown in pink) compared to the control (*Luc* shown in blue) are shown. Full statistics are shown in Supplemental Information. Related to S4, Table S10.

To infer regulators of clock differentiation we focused on the ISC population, since it is present at all three stages in intestinal development (Figure 3B, S4A, Table S10). We evaluated differences in regulon activity between the early pupa and immature adults to specifically address the changes occurring during these developmental stages as the clock first emerges (Figure 4B, Table S10). This analysis identified increased activity of 13 transcription factors, and decreased activity of 17 others during pupation. These include known ISC regulators *Stat92E,* and *GATA* (Buchon et al. 2009; Jiang et al. 2009; Beebe et al. 2010; Lin et al. 2010; Jiang and Edgar 2011; Okumura et al. 2016), the hormone signaling factors *gce*, *Met*, and *EcR* that are active during metamorphosis (Koelle et al. 1991; Riddiford and Truman 1993; Truman et al. 1994; Fisk and Thummel 1995; Lee et al. 2002; Li and White 2003; Abdou et al. 2011; Jindra et al. 2015; Zhang et al. 2021), and the clock gene *Pdp1*. Many other factors were identified that are specific to different cell types in the developing intestine. Our dataset provides a resource for the global transcriptional changes that occur during the growth and development of the *Drosophila* ISC model system.

*Drosophila* pupation consists of sequential hormonal pulses with rhythms in adult emergence occurring through circadian control of the ecdysone receptor (EcR) (Mark et al. 2021). We find that *EcR* and the nuclear receptor *ftz-f1* (*fushi tarazu transcription factor 1*), are downregulated in the immature adult ISCs, whereas *gce* (*germ cell-expressd bHLH-PAS*) and *met* (*Methoprene-tolerant*), both of which are involved in Juvenile hormone signalling during metamorphosis (Abdou et al. 2011; Jindra et al. 2015), are upregulated (Figure 4B-C). Some of these differentially expressed regulons also show clear bimodal distributions indicating that there are distinct cell populations, with and without activity of these transcription factors (*i.e. gce*, *Pdp1*, Figure 4C). This analysis provides a set of potential candidates that regulate the initiation of circadian clock activity.

To test whether these and other hormone nuclear receptors indeed regulate clock development, we performed a screen by depleting genes in a cell-specific manner. Using our scRNA-seq analysis as a guide, we depleted components of hormone signalling in ECs (*myo1A^TS^,* 23 genes) or ISC/EBs (*esg^TS^,* 10 genes) using RNAi in *Clock^TIM^*reporters during pupation (Fig 4D-E, S5C, Table S11). We reasoned that the loss of positive regulators of clock differentiation would delay Clk/cyc activity, while negative regulators would hasten it. We found that knockdown of components of ecdysone signalling in ECs (*EcR, usp, tai, Eip93F*) and Bursicon signaling (*rk*) decreased Clk/cyc activity on the first day after eclosion, compared to the control (Figure 4D). In ISC/EBs, we found that (as predicted by SCENIC) Clk/cyc activity was reduced by *gce* knockdown (Figure 4E). We also noticed that EC knockdown of *Eip93F* and *ftz-f1* fewer flies eclosed (Figure S5C) indicating an important developmental role. Of note, *Pdp1* loss (*Pdp1^3135^* mutant (Zheng et al. 2009)) did not affect initiation of Clk/cyc activity (Figure S5D), suggesting that this positive regulator of the circadian clock is not required for its development. These data are consistent with the notion that multiple hormone-signalling pathways in the intestine during pupal development regulate the emergence of the clock.

### The EC Lineage Reveals Heterogeneity in Clock Activity During Differentiation

EBs are progeny of ISCs that differentiate into ECs (Micchelli and Perrimon 2006; Ohlstein and Spradling 2006), but do not express clock genes robustly (Figure 3C). We therefore tested clock gene expression in the mature adult ISC to EC lineage. To distinguish between the steps in differentiation, we re-clustered and ordered adult cells from ISCs (cluster 0), EBs (clusters 2, and 3), and three EC populations (clusters 7, 8, 14) to reconstruct these stages in differentiation (Figure S5A). We then followed the longest lineage (ISC-0 to EC-7) to determine whether clock gene expression changes during EC differentiation (Figure 5A). The expression of a control housekeeping gene, *RpL32,* is present in all clusters, whereas the gene *klu* that correlates with the ISC identity (Korzelius et al. 2019; Hung et al. 2020b), and the EC marker gene *Amy-p* (Hung et al. 2020b) are enriched in their respective populations (Figure 5B). Clock gene expression was mapped on the same lineages and is reduced during the different stages of EB differentiation: *Pdp1, tim,* vri, and *cwo* are initially high in ISCs, then temporarily reduced in the EB populations, re-emerging in ECs when these complete differentiation (Figure 5B-C).

**Figure 5:**
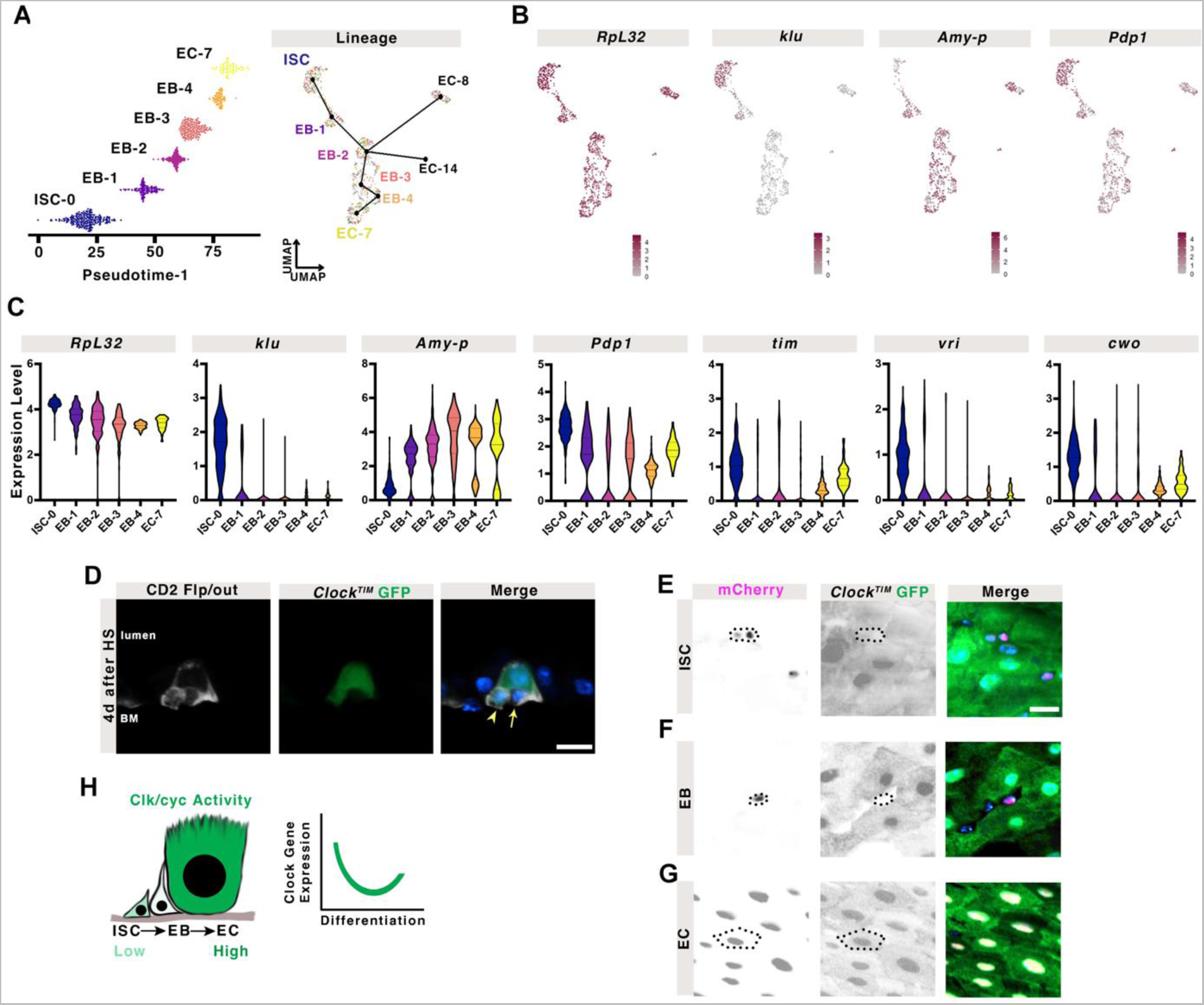
The EC Lineage Reveals Heterogeneity in Clock Activity During Differentiation. **(A)** A subset of clusters from the mature adult intestine (0,2,3,7,8,14) were assembled into a lineage from earliest cells (ISC = cluster 0) to differentiated ECs in pseudotime. See Figure S5A for more details. **(B)** Expression of a housekeeping gene, *RpL32*, shows it is expressed in all cell populations, whereas ISC-specific *klu* and EC-specific *Amy-p* are restricted to their respective populations. Mapping clock genes such as Pdp1 shows their expression in these lineage changes. **(C)** Graphs show cellular expression levels in this lineage of *RpL32*, compared to cell-specific genes (*klu* and *Amy-p*), and clock genes (*Pdp1*, *tim*, *vri*, *cwo*). *RpL32* is expressed throughout, *klu* only in ISCs, *Amy-p* only in differentiating EBs and ECs. Clock genes show initially high expression in mature ISCs which lowers in the transient differentiating EBs, to increase again in the EC. Lines indicate median and quartiles. Kruskal-Wallis test, full statistics are shown in Supplemental Information. **(D)** Flp/out clones show a mixture of basally located GFP+ and GFP- small cells, suggesting that Clk/cyc activity is heterogeneous in either ISCs and/or their progeny. Arrowhead marks GFP+ cell, arrow marks GFP- cell. DAPI marks nuclei, scale bar 10µm. Additional clones are shown in Figure S5B. Using mCherry expression to mark **(E)** ISCs, **(F)** EBs, and **(G)** ECs, specifically, Clk/cyc activity is present in ISCs, absent in EBs, and strongest in ECs. Outline indicates cell of interest marked with mCherry. DAPI marks nuclei, scale bar 10µm. **(H)** Schematic summarizing the observed decrease in Clk/cyc activity and clock gene expression during differentiation. Related to S5, Table S10-11.

These data indicate that clock genes are repressed in the ISC-to-EC lineage, hence we predicted that individual ISC clones should show Clk/cyc active and inactive cells. We used CD2-flp/out flies, where each CD2-marked clone consists of an initial marked ISC and all of its progeny, to lineage trace individual ISCs with a *Clock^TIM^* reporter. The clones revealed large GFP+ ECs, and a mixture of small GFP+ and small GFP-cells, consistent with the idea that a small GFP+ ISC produces small GFP-EBs that go on to differentiate into large polyploid GFP+ ECs (Figure 5D, S5B) (Parasram et al. 2018)). We further tested this idea by examining Clk/cyc activity specifically in only ISCs (*esg^TS^, Su(H)GBE-Gal80>mCherry*), only EBs (*Su(H)GBE>mCherry*), or only ECs (*Myo1A^TS^>mCherry*). We predicted that EBs should show lower Clk/cyc activity than their founder cells or their fully-differentiated progeny.

Indeed, Clk/cyc (*Clock^TIM^* reporter) activity is present in ISCs, absent in EBs, and highest in ECs (Figure 5E). These data are consistent with our scRNA-seq analysis and suggest the transient disappearance of circadian clock activity during lineage commitment of ISC to ECs in the adult intestine (Figure 4F).

### Photoperiod and Feeding Synchronize the Maturing Intestinal Circadian Clock

Our results indicate that the intestinal circadian clock gene expression is established during development, however, it is not clear what factors cause the intestine to exhibit synchronous circadian rhythms. The intestine completes its growth and development after adult emergence from pupation (O’Brien et al. 2011; Buchon et al. 2013), while it is being synchronized to the environment (Figure 2B-C). We have previously shown that the *Drosophila* intestinal clock is synchronized by photoperiod light cycles, with food ingestion also playing a modulating role in timing daily rhythms (Parasram et al. 2018). We first asked if photoperiod would be sufficient to synchronize the immature intestinal clock. Flies were completely starved for the first three days of adulthood following pupation and provided food only at ZT0 on adult day 4 when clock synchronization is complete (Figure 2B-C). Even though intestine size was reduced by lack of nutrients Clk/cyc activity was robust (Figure S6A), and synchronized rhythms emerged one day earlier (day 3 instead of day 4, compare Figure 6A to Figure 2B), suggesting that feeding and/or intestine growth delays clock entrainment.

**Figure 6:**
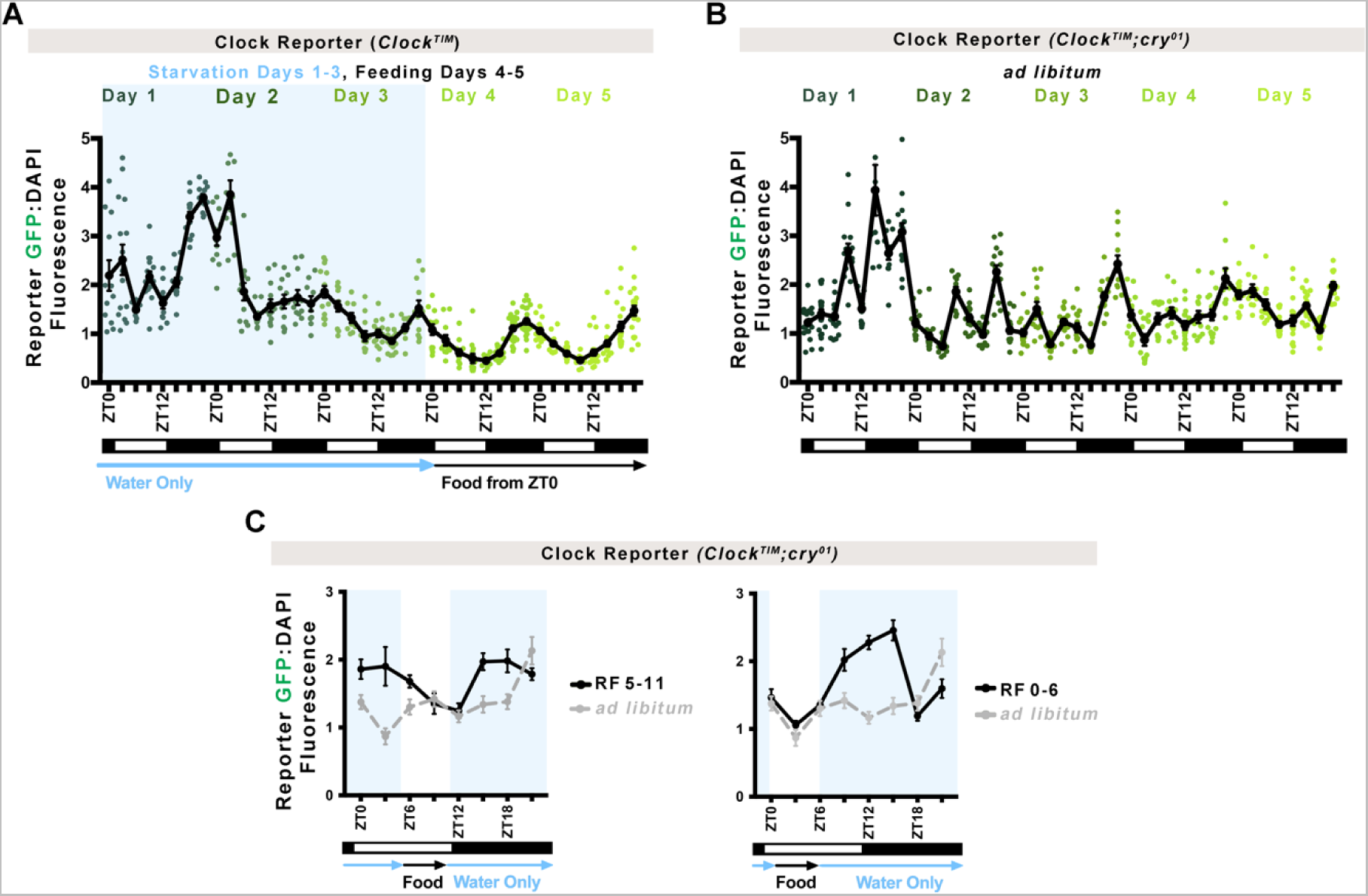
Photoperiod and Feeding Synchronize the Maturing Intestinal Circadian Clock. **(A)** *Clock^TIM^* flies that receive only water for the first three days after pupation show rhythmic Clk/cyc activity earlier: by day 3 rather than day 4, suggesting the absence of food more quickly synchronizes a rhythmic clock. Line shows mean. Error bars indicate ±SEM. Each point represents one intestine. One-Way ANOVA. **(B)** *Clock^TIM^;cry^01^*mutant flies that cannot sense photoperiod light display arrhythmic Clk/cyc transcriptional activity until day 4 when these are established, similar to controls. This suggests food in the absence of photoperiod can also synchronize a rhythmic clock. One-Way ANOVA. **(C,D)** Restricted feeding for 6 hours a day in *Clock^TIM^;cry^01^*mutants synchronizes intestinal Clk/cyc rhythms that peak ~6 hours after feeding. *Ad libitum* measurements (control) from Figure 4A are replotted in grey to show difference. Line shows mean. Error bars indicate ±SEM. Two-Way ANOVA. Full statistics are shown in Supplemental Information. Related to S6.

We then performed the opposite experiment, in which photoperiod is absent and feeding is present, by using *cry^01^* mutants that are unable to detect light to synchronize clock function (Dolezelova et al. 2007). We predicted that feeding would synchronize circadian rhythmicity. Without photoperiod, *cry^01^* intestines exhibit arrhythmic Clk/cyc activity during the first three days, but show Clk/cyc at day 4 onward similar to wildtype controls (Figure 6B compare to Figure 2B). Since light-responsive feeding behaviour occurs in *cry^01^* flies (Seay and Thummel 2011) these data suggest that feeding alone can also complete circadian clock synchronization in the intestine. To further test this, we restricted feeding in *cry^01^* flies post-pupation to the morning (ZT0-6) or evening (ZT5-11) and predicted that the timing of food consumption would alter the maxima and minima of intestinal circadian rhythms. Indeed, under these conditions the rhythms of *cry^01^* intestines were shifted to peak around 6h after feeding (Figure 6C). In contrast, control flies that have the ability to detect photoperiod do not show a shift in Clk/cyc activity on restricted feeding, suggesting that photoperiod is dominant over feeding if both environmental cues are detected (Figure S6B). We further tested if periods of fasting would affect clock rhythms once these are established in adults and noted that these do perturb the clock if photoperiod is detected (Figure S6C-D). We conclude that the combination of daily photoperiod and feeding cycles together synchronize the timing of intestinal circadian transcription cycles during clock maturation.

## Discussion

There are many questions about how circadian rhythms develop in the body (Vallone et al. 2007; Umemura and Yagita 2020) and, in particular, connections between circadian rhythms and stem cells are poorly understood (Brown 2014; Benitah and Welz 2020). We show that as the intestine grows and differentiates clock gene transcription is negatively regulated (Figures 1, 3, 4); ISCs differentiating into EB precursors also do not express clock genes (Fig 5). Our study sheds light on two processes: 1) the emergence of circadian clock function in the stem cells of the intestine and 2) suppression in their immediate progeny as these establish a differentiated cell state. These data suggest that the clock and differentiation are incompatible in the intestine and are reminiscent of a recent report showing clock function is incompatible with mouse somite development (Umemura et al. 2022). Our study characterizes the development of the circadian clock in the intestine but this framework may be extended to other organs of the body.

Circadian clock development has been tested in several animal species. In zebrafish, clocks emerge during the first stages of embryogenesis (Dekens and Whitmore 2008), similarly, in *Drosophila* clock emergence was shown to occur early in development (Sehgal et al. 1992). However, these initial studies of *Drosophila* clock development focused on circadian rhythms in the brain rather than other tissues in the body (Sehgal et al. 1992; Keene et al. 2011). It was recently found that *Drosophila* larval tissues do not express the clock gene *cry* until adulthood (Agrawal et al. 2017), and circadian control of the prothoracic gland also occurs in pupation (Di Cara and King-Jones 2016), consistent with our results (Figure 1). Of note, we find that in Clk/cyc activity is initiated in the spiracles and prothoracic gland during larval development, and the heart during pupation (unpublished observations). This suggests that circadian timing in the *Drosophila* central clock emerges earlier and timing in other tissues emerges later. In chicks, even though clock expression occurs early in development, it is patchy during early stages before becoming more widely expressed (Gonçalves et al. 2012). In mice and rats, where circadian development is best studied, early embryonic cells do not have clock rhythms (Kowalska et al. 2010; Yagita et al. 2010). The central pacemaker in the brain emerges during late embryogenesis (Sládek et al. 2004; Landgraf et al. 2014; Carmona-Alcocer et al. 2018), and other tissues such as the heart (Umemura et al. 2017), kidney (Dan et al. 2020), and colon (Polidarová et al. 2014) emerge even later. These mammalian tissue clocks are present just before birth and their synchronous rhythms mature at different rates postnatally (Yamazaki et al. 2009), particularly in the colon that (like our study) shows the intestine has a relatively slow clock to develop in the body (Polidarová et al. 2014). Overall, data from a wide variety of species are consistent with our observations. Circadian clock function is also tissue-specific, and thus its emergence may be tissue-specific, eventually coalescing into complete circadian rhythmicity throughout the body at the appropriate stage of development. The larval stages where insects eat almost continuously may be incompatible with a daily timing system in the digestive tract that dictates activity/rest cycles.

Our study is the first to examine the emergence of animal tissue clocks at the single cell level, revealing that stem cells are the first to pass clock status to their differentiating progeny. The transient loss of clock gene expression as cells transition between discrete fates has been previously overlooked when using population-based methods such as RNA-seq or RT-qPCR. Thus, our study provides a paradigm of how clocks may develop in non-neural tissue cells after the central clock is established. We have used the expression of several clock components simultaneously to estimate the developmental timing of clock rhythms. It is important to recognize that the expression of clock genes alone does not confirm such rhythms are present. Indeed, we find that clock gene expression is present in the early adult intestine but is not rhythmic until several days later (Figure 1-2). The early clock in the zebrafish circadian system also takes 3-4 days to be established, with the secondary feedback system and output genes rhythmic later in development (Ziv and Gothilf 2006; Laranjeiro and Whitmore 2014). This 2-4 day delay between gene expression and daily cycling also appears even in the mouse kidney (Dan et al. 2020) and central clock in the brain (Carmona-Alcocer et al. 2018).

Elegant studies in mouse embryonic cells have shown that translational regulation of *Clock* (the mouse ortholog of *Clk*) is a rate-limiting step in circadian development (Umemura et al. 2017). Our results are in agreement with this (Figure 1H), however, in *Drosophila* we identify the component *Pdp1* as another potential factor (Figure 1H, 3C, S4C). This is consistent with a Clk-based transcriptional model because Pdp1 positively regulates *Clk* expression, hence *Clk* expression and transcriptional activity may be a conserved circadian component that is required for rhythmicity. Although loss of *Pdp1* does not affect the timing of intestinal clock development, in *Drosophila*, it has been shown that ectopic expression of *Clk* can prematurely induce clocks outside the pacemaker (Zhao et al. 2003; Kilman and Allada 2009). Future work to characterize this potential mechanism in ISCs will be informative, as it may be the switch that initiates daily tissue timing in developing tissues at different times.

To our knowledge, this is the first single-cell characterization of insect intestinal metamorphosis and circadian clock development. Because insects utilize circadian rhythms to coordinate their pupation timing (Di Cara and King-Jones 2013), these results are informative in understanding the fundamental biology of these animals. Our analysis with SCENIC (Aibar et al. 2017) identified key transcription factors involved in the development of this organ, showing the existence of transitory Pupal Cells (PupCs) that express markers of early differentiation such as *Sox14* (Meng and Biteau 2015). Using scRNA-seq, we were able to recover the main cell types in the intestine at all three stages and found markers to identify ISCs, EBs, EEs, ECs, and visceral muscle (VM) that are consistent with previous *Drosophila* intestine sequencing studies (Buchon et al. 2013; Marianes and Spradling 2013; Dutta et al. 2015; Hung et al. 2020b; Li et al. 2022) (Table S10). We were not able to identify a population of enteroendocrine mother cells although the earliest ISC population includes cells enriched for *piezo*, a marker of the EE lineage (He et al. 2018), therefore it is likely that these cells are located in the ISC cluster (Cluster 0, Figure 3A). We have annotated a population of pupal cells, PupCs, that are unique to the pupal intestine and display similarities to differentiated cells including expression of *MtnB* and *MtnD* characteristic of the central intestine, Table S10) suggesting that these cells contribute to intestine development consistent with the transient pupal midgut described previously (Takashima et al. 2011b). Our data provides a resource for the development of the intestine in insects during metamorphosis.

By testing clock development at the single cell level, we found that the circadian clock program is downregulated during the dynamic cellular changes that occur during differentiation. These results are counterintuitive given that the clock is thought to emerge during embryogenesis, and has in fact been demonstrated to arise in differentiating embryonic stem cells (Landgraf et al. 2014; Carmona-Alcocer et al. 2018), and organoids (Rosselot et al. 2022) *in vitro*. Indeed, several studies have shown circadian clocks regulate cellular differentiation itself (Yu et al. 2013; Alvarez-Dominguez et al. 2020; Zhang et al. 2022). We therefore propose that during transient states of cellular differentiation, clock gene expression is disrupted. A parsimonious explanation for this is that the transcriptional and epigenetic reorganization that occurs during differentiation (Jin et al. 2013; Ma et al. 2013; Boumard and Bardin 2021) simply interferes with rhythmic circadian transcription. Interestingly, a recent study observed a similar pattern of circadian disruption in thymocyte differentiation where a Per1-Venus reporter exhibits reduction in cellular stages between thymocyte precursor and fully differentiated cell (Hemmers and Rudensky 2015). In line with our data, this suggests that clock function requires a certain level of epigenetic stability that is not compatible with the rapid cellular fate changes that occur during development or during stem cell-driven regeneration. Our system provides an opportunity for future investigation of these interactions and highlight a state where cells in the body are resistant to circadian rhythms. Because both ISC differentiation (this study) and somite differentiation (Umemura et al. 2022) involve cell fate commitment through Notch signaling, it will be of interest to see if these principles extend to other developmental pathways and/or confirm the precise mechanisms linking the clock and the Notch pathway in the future.

In human beings, circadian rhythmicity arises postnatally: most infants take several months to establish coherent daily timing (Rivkees 2003). This is likely to reflect the complexity of establishing robust inter-cellular and inter-organ synchrony that involve coordinating environmental signals with tissue-specific clocks. Our study reveals how circadian rhythms are established in complex developing tissues and provides a system in which to investigate the birth of the circadian tissue pacemaker.

## Materials and Methods

### Fly Strains

*Clock^PER^, Clock^TIM^* described in [1]. From Jadwiga Giebultowicz: *CantonS* and *per^01^*(isogenic with *CantonS*). From Chrysoula Pitsouli *esg-Gal4 (II), tubGal80^TS^(III).* From Yong Zhang *PER-AID-GFP*. From Steve Jean *Su(H)GBE-Gal80, tubGal80ts(III).* From Amita Sehgal *Pdp1^3135^(III).* From Bloomington: #38424 *UAS-mCherry (III), UAS-mCherry(II), myo1A-Gal4 (II),* TRiP #JF01355 *Valium1-Luc(III),* #42563 *Valium20-cyc(II), cry^01^*, #76317 *CRY-GFP, #36634 Blimp-1 RNAi, #4412 act>y+>Gal4,UAS-CD2, RNAi lines: #36634 Blimp-1, #29373 and #41707 dsf, #50712 and #58286 EcR (#1 and #2), #43231 Eip75B, #26718 Eip78C, #57868 Eip93F, #61937 and #50868 ERR, #27659 ftz-f1, #26323 and #61852 gce, #29375 and #64988 Hnf4, #51442 and #27254 Hr3 (#1 and #2), #29376 and #29377 Hr38, #33624 and #27086 Hr39, #54803 and #31868 Hr4, #39032 Hr51, #31990 Hr78, #31990 Hr96, #31603 Luciferase, #26205 met, #31958 rk, #28689 svp, #36095 tai, #27242 tll, #36729 and #27258 Usp (#1 and #2)*. Flies were balanced with *yw;IF/CyO;MKRS/TM6B*. Flies were raised on cornmeal-glucose media (1.2% w/v yeast, 0.7% w/v soyflour, 5% w/v cornmeal, 0.4% w/v malt, 0.4% v/v agar, 5.3% v/v glucose with propionic acid and tegosept) at 25°C with 12:12 light/dark photoperiod (lights-on at ZT0). For free-running experiments flies were shifted to constant darkness (DD) during the dark-phase of the final day of LD, CT0 (Circadian Time 0) is the time when lights are expected to turn on. For larval to adult analysis, flies were only allowed to lay eggs for 5 hours in the morning (ZT0-5). For adult experiments, only flies that eclosed overnight (ZT12-0) were included, no difference was noted between ZT11-12, ZT23-0, and ZT0-1 eclosion groups. Heatshocks for Flp/out clones were done with adult flies 5-14 days old for 5-10min at 37°C. For RNAi screening (Figure 4D-E), flies were raised at 18°C until pupal stages then vials were shifted to 29°C and both males and females were collected.

### Food Restriction

Pupa were transferred to vials containing a piece of water-moistened filter paper using a paintbrush. Flies were kept on water after eclosion either (1) until adult day 4 at ZT0 when flies were transferred to vials containing normal food or (2) for 6 hours (from either ZT5-11 or ZT0-5) each day after eclosion.

### Developmental Staging

Larva were roughly staged using the following times: 1 day AED (after egg deposition) for L1, 2 days AED is L2, 3-5 days AED is L3 (the day AED may be noted e.g., L3 day 3 refers to three days AED). Late stage pupae were analyzed at the grey to black wing stage (day 9 or 10 AED) [2]. The first day after eclosion is referred to as adult day 1 (day 9 or 10 AED).

### Dissection

Flies were euthanized in 70% EtOH and then intestines were dissected using fine forceps in 1xPBS. For RNA extractions and single cell analysis, the hindgut (anterior to the Malpighian tubules) and the foregut were removed during dissection to ensure only the intestine was sampled. Dissections were completed in <1 hour. Analysis of scRNA-seq from larva to adult included both males and females but unless otherwise noted, only female flies were analyzed.

### Tissue Preparation / Antibody Staining

Intestines were fixed in 4% paraformaldehyde for 40 minutes, washed twice in 1xPBS and stained with DAPI (1:5000 in 0.2% Triton-X100 (in 1xPBS)) for 5 minutes. The intestines were washed twice in 0.2% PBS-T and then mounted on slides using AlexaProLong Gold Antifade Reagent (Invitrogen). For antibody staining, intestines were fixed for 2 hours in 4% paraformaldehyde and then washed twice in 1xPBS and blocked for 20 minutes in 1% bovine serum albumin in 0.2% PBS-T. The intestines were then incubated in primary antibody (PER (1:1200), histone (1:2000), GFP (1:2000), rat-CD2 (1:2000)) in BSA for 2 hours, washed twice in 0.2% PBS-T, incubated in secondary antibody (1:2000) with DAPI (1:5000) in BSA for 1 hour and then washed twice in 0.2% PBS-T (at 4°C) and mounted on slides.

### RT-qPCR

Approximately 15 female flies of *CantonS* were dissected in 1XPBS before being transferred to tubes containing RNAlater (Qiagen) on ice. RNA was extracted using the RNAMini Kit (Qiagen) with RLT buffer (Qiagen) using a Bullet Blender (Next Advance) for homogenization. cDNA was made using ISCRIPT RT Supermix (Bio-Rad) and RT-qPCR was performed using SYBR green (Bio-Rad) using the Viia7 PCR plate reader, primers: per-F TCATCCAGAACGGTTGCTACG, per-R CCTGAAAGACGCGATGGTGT [3], tim-F CCAGCATTCATTCCAAGCAG, tim-R GCGTGGCAAACTGTGTTATG [3], Gapdh-F CCAATGTCTCCGTTGTGGA, Gadph-R TCGGTGTAGCCCAGGATT, Clk-F CAAGTTTGGCCTCTGGCTCTC, Clk-R TACAACTAGCTCTGGGCTTCCG, Pdp1-F GAACCCAAGTGTAAAGACAATGCG, Pdp1-R CTGGAAATACTGCGACAATGTGG [4], vri-F TGTTTTTTGCCGCTTCGGTCA, vri-R TTACGACACCAAACGATCGA [4], cyc-F GCGCTGATGGAGTCTCACAAG, cyc-R GTAGCTGTTGTCCTTGCACCG, cry-F CACCGCTGACCTACCAAA, cry-R GGTGGAAGCCCAATAATTTGC [5], dco-F TTGGGAGGAGGGTTAGCAG, dco-R TTACAATGTGGGTGCCTTGC.

### scRNA-seq

Flies from *CantonS* were euthanized in 70% ethanol in DEPC-treated water at ZT0. Every 10 minutes during the 1h dissection time, the intestines were transferred from the 1XPBS well to ice-cold 1% BSA. Once dissections were complete, the intestines were transferred to a drop of 1XPBS on the back of a dissection plate and cut into small pieces using microscissors. The fragments were then transferred to a 1.5mL tube to a total volume of 350ul (200uL for pupa). Elastase was added to a concentration of 1mg/mL and incubated at 37°C for 40min pipetting up and down 30-40 times every 10 minutes to dissociate the cells, then 10% BSA was added to a final concentration of 1% BSA. The samples were centrifuged at 300xg for 15min at 4°C, then the pellet was resuspended in 200ul of 0.04% BSA and filtered with a 70um filter centrifuging at 300xg at 4°C for 1min. Orbitrap Density Gradient was used to sort the cells and then the interface was washed with 0.04% BSA. The cell density was determined with a haemocytometer and diluted with 0.04% BSA to a density of 800cells/uL. All samples had a viability >95%. The entire preparation was completed within 2 hours. The fresh samples were submitted to the High Content Analysis Core at the University of Alberta by ZT3 for 10X Genomics 3’ Library preparation and to Novogene for sequencing (NovaSeq) and then aligned to the Drosophila reference genome 6.25 using Cell Ranger.

### Imaging and Data Analysis

Slides were imaged with either a slide scanner (Zeiss, Axio Scan Z.1) or confocal microscope (Olympus, IX81 FV1000 with 60x Water Immersion Lens or Zeiss Confocal with 20x lens). Whole mount imaging and quantification were done using a fluorescence microscope (Zeiss, VertA.1) or fluorescence stereoscope (Leica M205). Images were analyzed with Zen Blue Software (Zeiss) and processed using Photoshop (Adobe). Fluorescence intensity measurements are ratios of GFP:DAPI. RT-qPCR data is normalized to *Gapdh1*. Prism (GraphPad) was used for graphing and statistical analysis.

Single sequencing data was analysed in R using Seurat and pseudotime analysis was performed using Slingshot. The original dataset of 6835 adult, 2625 pharate, 3816 pupal cells was filtered (genes must be detected in >=3 cells, %mitochondrial <25, 200<nFeatures<4000, 1000<nCount<100000) and the remaining 1980 adult, 1197 pharate, 2013 pupal cells were used for further analysis. The datasets were integrated (reduction=rpca) and clustered (PCs=14, resolution=0.5) and then annotated using differentially expressed genes (method=roc). For lineage tracing 1111 adult cells consisting of clusters 0,2,3,7,8,14 were re-clustered (PCs=9, resolution=0.3) or 1933 pupal cells from clusters (0-6,8-14,16,17,18,21) were re-clustered (PCs=16, resolution=0.3) prior to lineage tracing to further subdivide these cells. Differential gene analysis was performed using roc. SCENIC analysis was performed with version 10 *Drosophila* motif set with minimum number of genes in each regulon=10, then handed back to Seurat for differential gene expression anlaysis. In some cases, Prism (GraphPad) was used for graphing and statistical analysis. Images were processed using Photoshop (Adobe). scRNA-seq data have been deposited at GEO (GSE230572) and are publicly available as of the date of publication.

Statistical tests were performed using GraphPad Prism and can be found in Supplemental Information.

## Abbreviations

ISC: intestinal stem cell
EB: enteroblast
EC: enterocyte
EE: enteroendocrine cell
PupC: pupal cell
AMP: adult midgut precursor
PC: peripheral cells
scRNA-seq: single cell ribonucleic acid sequencing
Myo1A: myosin 31DF
esg: escargot
per: period
tim: timeless
Clk: *Drosophila* circadian locomotor output cycles kaput or clock
cyc: cycle
Pdp1: par domain protein-1
vri: vrille
dbt: doubletime also known as dco – discs overgrown
cry: cryptochrome
cwo: clockwork orange
klu: klumpfuss
Amy-p: amylase p
RpL32: ribosomal protein L32
EcR: ecdysone receptor
Eip93F: Ecdysone-induced protein at 93F
gce: germ cell-expressed bHLH-PAS
met: Methoprene-tolerant
ftz-f1: fushi tarazu transcription factor 1
usp: ultraspiracle
ERR: estrogen-related receptor
GFP: green fluorescent protein
AID: auxin inducible degradation
CantonS: Canton (Special)

## Acknowledgments

We are grateful to the members of Karpowicz and Foley labs and the technical staff at the Universities of Windsor and Alberta for their insightful advice and expertise. KP, AZ, and PK were funded by the Canada Foundation for Innovation, the Ontario Research Fund, and the Natural Sciences and Engineering Research Council of Canada. BD was funded by the Natural Sciences and Engineering Research Council of Canada. MS, RW, and EF were funded by Canadian Institutes of Health Research. To the authors whose research we could not cite due to space limitations, we offer our apologies and thanks.

## Author Contributions

Investigation, KP, AZ, MS, and RW; Formal Analysis, KP, AZ, RW, BD; Writing – Original Draft, KP and PK; Writing – Review & Editing, KP, BD, EF, and PK; Conceptualization, PK; Supervision, EF and PK; Funding Acquisition, EF and PK

